# Ancient Y chromosomes confirm origin of modern human paternal lineages in Asia rather than Africa

**DOI:** 10.1101/2020.03.10.986042

**Authors:** Hongyao Chen, Ye Zhang, Shi Huang

**Affiliations:** Center for Medical Genetics, Hunan Key Laboratory of Medical Genetics, School of Life Sciences, Central South University, 110 Xiangya Road, Changsha, Hunan 410078, P.R. China

**Keywords:** Y chromosomes, ancient DNAs, Out of East Asia, Out of Africa, infinite site model

## Abstract

Analyses of Y chromosome variations of extant people have resulted in two models for the paternal phylogenetic tree of modern humans with roots either in Africa or East Asia. These two trees are differentiated mainly by when and where their mega-haplogroups branched apart. This paper examines previously published Y chromosome sequencing data of 17 ancient samples to compare these two competing models. As ancient samples have had less time to evolve, they are expected to have mutated in some, but not all, of the sites that define present day haplogroups to which they belong. Indeed, most of the ancient DNAs here showed that expected pattern for both the terminal and the basal haplogroups to which they belong, all of the ones which are non-controversial or considered real by both of the two competing models followed that pattern. However, for basal haplogroups not shared by the two models, such expected pattern could be observed only if the haplogroups specific to the Asia rather than the Africa model are real, including ABCDE, ABDE, AB, A00-A1b. Another important point is that, if the mega-haplogroups of the Africa model were real, including BT, CT, CF and F, it would mean that numerous alleles would be shared between these haplogroups and several ancient A1b1b2 samples, which is unexpected and unseen in present day samples. Sharing alleles like this would also violate the infinite site assumption that makes the Africa rooting possible in the first place. Therefore, the data from ancient Y chromosomes confirm the actual existence of the haplogroups specific to the Asia model.

---

Two competing models of modern human origins termed “Multiregional” and “Recent Out-of-Africa” have long been accepted ^1,2^. The multiregional model considers extant people of any given region, e.g., East Asia, to be largely descended from ancient people living in the same region at ~200-2000 ky ago, such as Peking man. The model has support from fossils and cultural remains but molecular evidence has been lacking until recently ^3^. Analyses based on a more complete molecular evolutionary framework, the maximum genetic diversity (MGD) theory ^4^, suggest multiregional origins for autosomes but root both uniparental DNAs in East Asia ^3^. The rooting of mtDNA tree in Asia independently confirms an earlier paper ^5^, and has been verified by ancient mtDNAs findings that show the earlier appearance of haplogroup R compared to N ^6,7^. The Out of Africa model posits that modern humans originated in Africa and then migrated to Eurasia, largely replacing local archaic humans with limited genetic mixing ^1,8,9^. The rooting of uniparental DNAs in Africa relies on the assumption of neutral mutations throughout the entire genome ^9–11^, which is known to be a poor explanatory framework for evolutionary phenomena ^4,12^. The infinite site assumption emerges from the neutral framework, which states that mutations appear once in the evolutionary history, and the related inference of derived alleles underlies the Y tree topology and rooting of the Africa model ^10^. However, mutation saturation and natural selection is far more common than initially thought, which would invalidate the currently accepted inference of derived alleles ^3,4,13,14^. Certain haplogroups contain a large number of derived alleles that define other haplogroups, e.g., A has many derived alleles for BT (42.4% of informative sites) and A and B have many derived alleles for CT (18.9% informative sites), which violates the method underlying the Africa model in the first place ^3,15^. In contrast, haplogroups in the Asia model are defined by alleles shared by all members within a haplogroup, regardless of their derived status. The rooting in the Asia model of uniparental DNAs relies on the reasoning that the original haplotype should be the common type, since mutations leading to alternative types should be rare events ^3,5–7^. The ancestor type should have many alleles different from the outgroup to qualify as a modern type. Some of those alleles may revert back to archaic alleles as modern types migrated to new environments and admixed with archaic humans. Co-evolution with admixed autosomes may cause modern uniparental DNAs to mutate back to archaic alleles.

The biggest difference between the two competing models of Y tree topology comes from how they handle the mega-haplogroups in the Africa model upstream of G (Figure 1), such as BT or CT. This category includes the vast majority of haplotypes among present day diversity and is thought to result from new mutations since the original type. In contrast, the Asia model considers the alleles of mega-haplogroups like BT or CT to be the ancestor type carried by the first modern human individual. It is the non BT or non CT haplotypes such as A or AB that have acquired new mutations. We here used ancient DNAs to test these two competing models.

**Figure 1.**
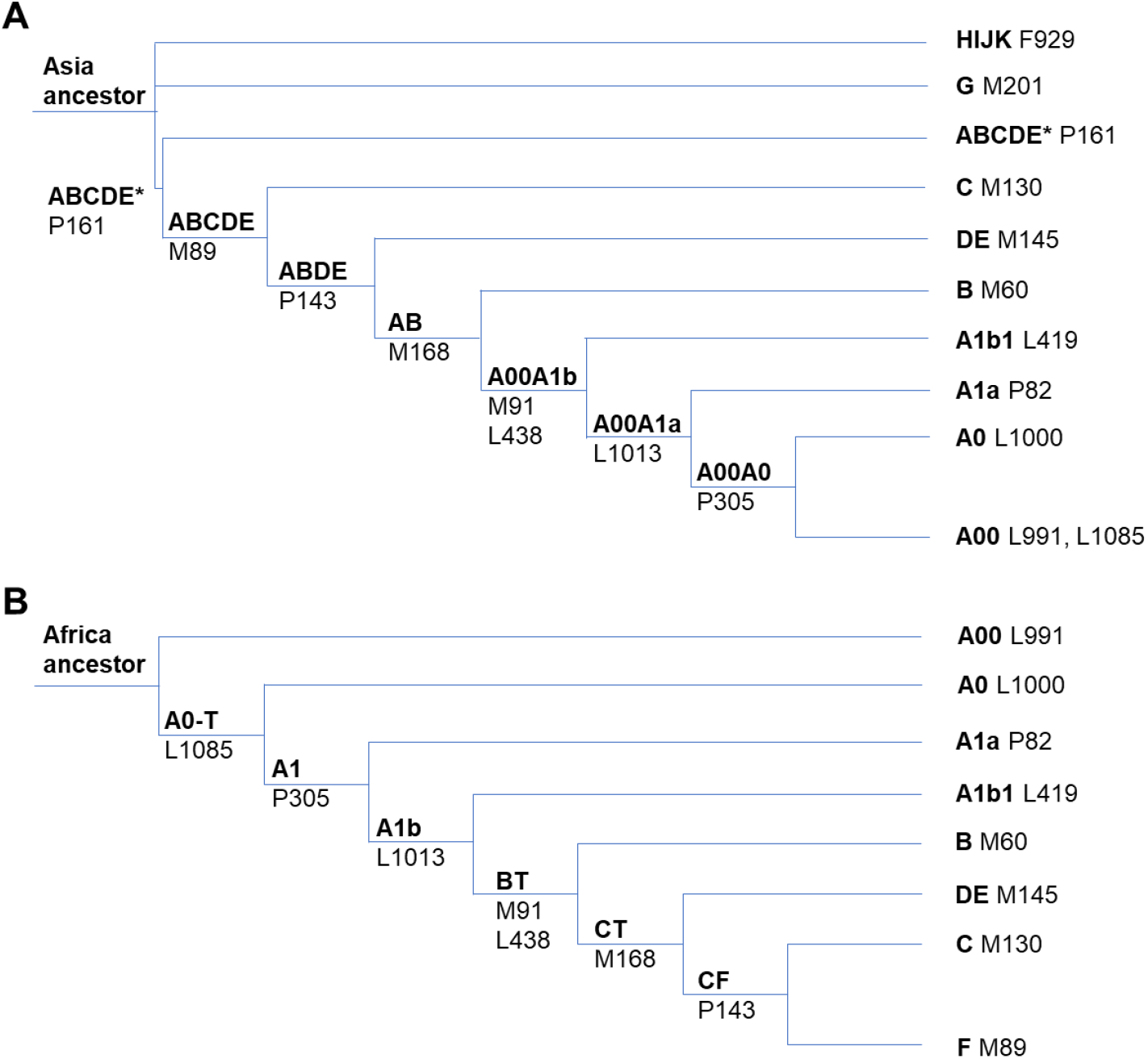
Y chromosome phylogenetic trees of modern humans. Only major branches and representative SNPs are shown with branch lengths not to scale. The tree topology was built without making use of any ancient DNAs. A. The Out of East Asia model. B. The Out of Africa model.

To confirm the expectation that ancient samples should have mutated in only a fraction of the sites that define the present day haplogroups to which they belong, we studied a total of 17 published ancient Y chromosome sequences of relatively high coverage (Table 1), including A1b1, B, C, E, H, I, and R haplogroups ^16–23^, for informative sites as found in the 1000 genomes project (1kGP) or in the Y-DNA haplogroup tree from the International Society of Genetic Genealogy (ISOGG, http://www.isogg.org, version 13.07) ^24^. Most of these ancient DNAs showed mutations in some but not all of the sites that define the present day haplotypes to which they belong (Figure 2A, Supplementary Table S1 and S2). For example, ATP12-1240 sample from Atapuerca Spain had 58 out of 67 informative sites mutated for I-M170 and 20 of 25 informative sites mutated for the basal haplogroup IJ-M429 ^21^; I2966 sample from Malawi had 3 of 3 informative sites mutated for B2b1, 30 of 37 informative sites mutated for B2-M182 and 7 of 9 mutated for the basal branch B-M181 ^16^. To exclude the possibility of sequencing or calling errors, we studied all samples for mutations in sites that define an irrelevant haplogroup A1a and found all to have essentially no mutations in A1a, as expected if there were essentially no calling errors (Figure 2B, Table 1). Only an extremely low rate of unexpected changes was found (9 out of 9882 A1a sites called for all samples), which may be either mutations or sequencing/calling errors.

**Figure 2.**
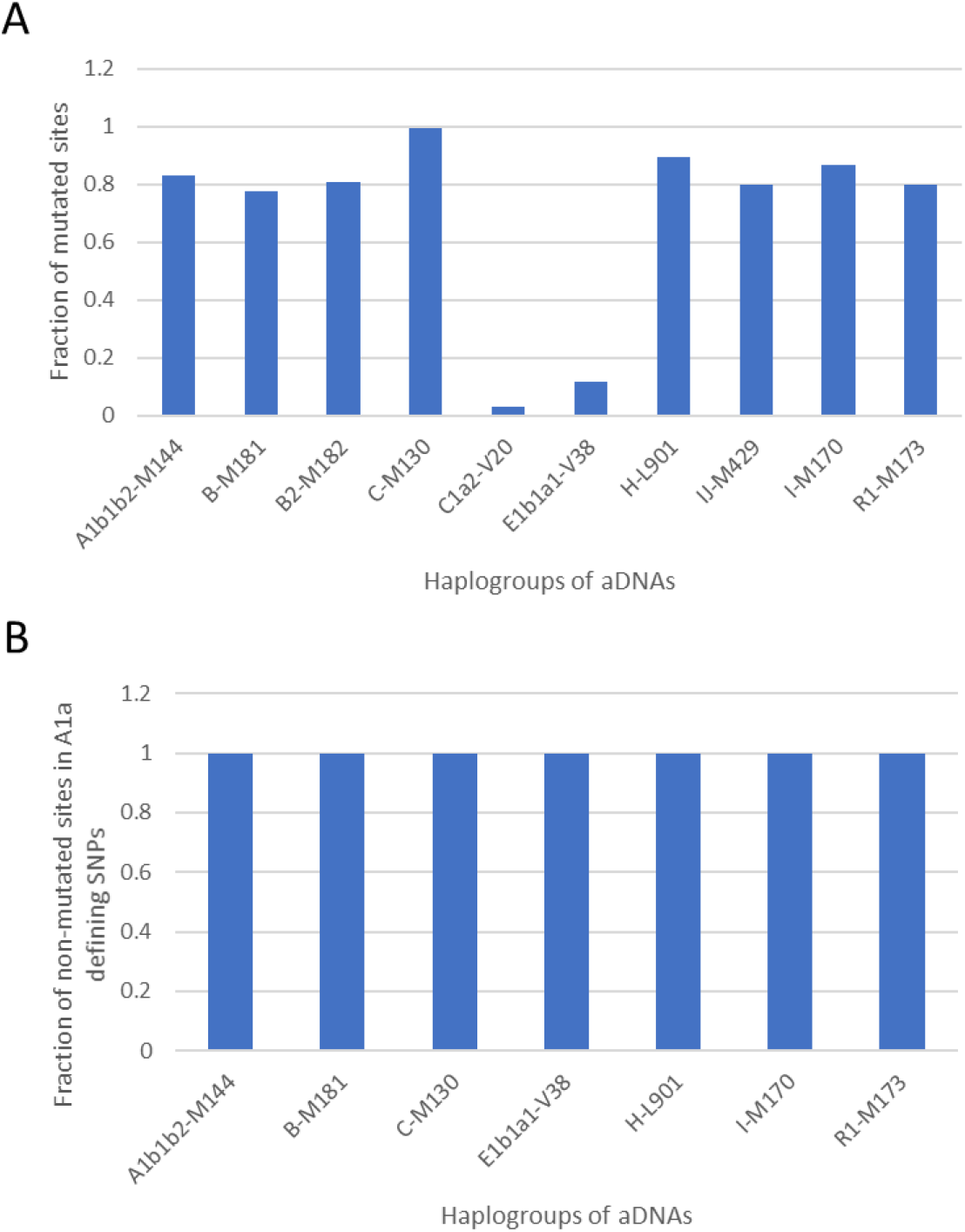
Incomplete mutations in ancient Y chromosomes in haplogroups regarded as real by both the Africa and Asia models. A. The fractions of mutated sites among the informative sites defining a haplogroup were shown for ancient samples belonging to each haplogroup as shown. B. The fractions of non-mutated sites among the informative sites defining A1a haplogroup were shown for ancient samples belonging to each haplogroup as shown. Most of the listed haplogroups have only one ancient sample but some have more than one.

**Table 1.** Mutation patterns in ancient Y chromosomes. Haplogroup-defining SNPs were identified using ISOGG and the 1kGP dataset. The first number in the site # row refers to the number of haplotype-defining sites in ISOGG and the second number refers to sites found in the 1kGP. The total combined number of sites from ISOGG and 1kGP was used in the analysis. For numbers in the remaining cells, the first number refers to mutations in haplotypes of the Out of East Asia (OOEA) model, the second number refers to mutations in haplotypes of the Out of Africa (OOA) model, and the third number represents the total number of informative sites. Numbers in bold highlight cases where only a fraction of the sites defining a basal haplogroup were mutated.

| Ancient samples | Year BP | OOEA | A00A0 | A1a | A00A1a | A00A1b | AB | ABDE | ABCDE |
| --- | --- | --- | --- | --- | --- | --- | --- | --- | --- |
|  |  | OOA | A1 | A1a | A1b | BT | CT | CF | F |
|  |  | HG/Site# | 12, 0 | 13, 936 | 35, 4 | 314, 0 | 177, 0 | 3, 1 | 58, 93 |
| Ballito Bay A, S Afr | 1986 | A1b1b2 | 0/12/12 | 1/1/946 | 0/39/39 | **302/9/311** | **171/2/173** | 4/0/4 | **144/5/149** |
| Ballito Bay B, S Afr. | 2149 | A1b1b2 | 0/9/9 | 1/1/466 | 0/19/19 | **131/24/155** | **66/12/78** | 2/0/2 | **56/13/69** |
| I9028, S Afr | 2000 | A1b1b2a | 0/4/4 | 2/2/422 | 0/22/22 | **77/52/129** | **41/37/78** | 0/0/0 | **37/23/60** |
| I9133, S Afr | 2000 | A1b1b2a | 0/7/7 | 0/0/647 | 0/24/24 | **128/55/183** | **85/27/112** | **1/1/2** | **63/22/85** |
| I2966, Malawi | 8100 | B2b1 | 0/3/3 | 0/0/91 | 0/7/7 | **1/91/92** | **32/18/50** | 2/0/2 | **22/9/31** |
| Mota, Ethiopia | 4500 | E1b | 0/11/11 | 0/0/899 | 0/38/38 | 0/301/301 | 0/172/172 | 4/0/4 | **117/2/119** |
| K14, Russia | 36000 | C1b | 0/7/7 | 0/0/355 | 0/14/14 | 0/109/109 | 0/70/70 | 0/1/1 | **15/3/18** |
| Sunghur SI, Russia | 36000 | C1a2 | 0/3/3 | 0/0/361 | 0/17/17 | 0/148/148 | 0/88/88 | 0/3/3 | 16/0/16 |
| Sunghur SII, Russia | 36000 | C1a2 | 0/11/11 | 0/0/809 | 0/36/36 | 0/278/278 | 0/155/155 | 0/4/4 | 87/0/87 |
| Sunghur SIII, Russia | 36000 | C1a2 | 0/12/12 | 0/0/925 | 0/39/39 | 0/310/310 | 0/174/174 | 0/4/4 | 139/0/139 |
| Sunghur SIV, Russia | 36000 | C1a2 | 0/11/11 | 0/0/799 | 0/35/35 | 0/265/265 | 0/153/153 | 0/4/4 | 85/0/85 |
| Vestonice 16, Czech | 30000 | C1a2 | 0/5/5 | 0/0/83 | 0/6/6 | 0/87/87 | 0/59/59 | 0/2/2 | **18/1/19** |
| Brana 1, Spain | 7500 | C1a2 | 0/7/7 | 0/0/672 | 0/28/28 | 0/230/230 | 0/127/127 | 0/4/4 | 46/0/46 |
| ATP2, Spain | 4900 | H | 0/11/11 | 1/1/869 | 0/37/37 | 0/282/282 | 0/162/162 | 0/4/4 | 0/127/127 |
| ATP12-1420, Spain | 4900 | I2 | 0/7/7 | 4/4/482 | 0/17/17 | 0/164/164 | 0/95/95 | 0/2/2 | 0/70/70 |
| MA1, Russia | 24000 | R1-M173 | 0/8/8 | 0/0/465 | 0/24/24 | 0/167/167 | 0/88/88 | 0/1/1 | 0/78/78 |
| F38, Zagros | 4500 | R1b-M343 | 0/9/9 | 0/0/591 | 0/23/23 | 0/204/204 | 0/130/130 | 0/4/4 | 0/92/92 |

Many but not all ancient DNAs showed the expected mutation pattern for basal branches specific to the Asia model such as A (A00-A1b), AB, ABDE, and ABCDE, and they all showed a near complete absence of mutations in the haplogroups to which they do not belong (Figure 3A and B, Table 1). For example, for the five samples belonging to the AB haplogroup, there were a total of 395 mutations among 491 informative sites that classify them as AB while the remaining 96 sites are shared with non AB haplogroups and represent the genotype of the original type. In contrast, essentially no ancient DNAs showed the expected mutation pattern for branches specific to the Africa model such as A1, A1b, BT, CT, CF, and F, and some showed unexpected mutations in the haplogroups to which they do not belong (Figure 3C and D, Table 1). For example, for the 12 samples belonging to the CT haplogroup, all 1473 informative sites would have mutated to CT-specific alleles if the CT branch was legitimate. There was only one case with the expected pattern, if any of the mega-haplogroup in the Africa model are real, where the I2966 B2b1 sample had 91 of 92 sites mutated to BT (Table 1). However, as this involved only 1 out of 92 sites, it may just be a sequencing/calling error. In addition, the five non-CT samples had 96 out of 491 informative sites for CT mutated to CT-specific alleles. As no A or B haplogroup today are known to share CT alleles in CT defining sites, the finding of ancient A or B samples carrying CT alleles is highly unusual and unexpected. It would mean that those sites have mutated more than once, which would violate the infinite site assumption that makes the Africa rooting possible in the first place.

**Figure 3.**
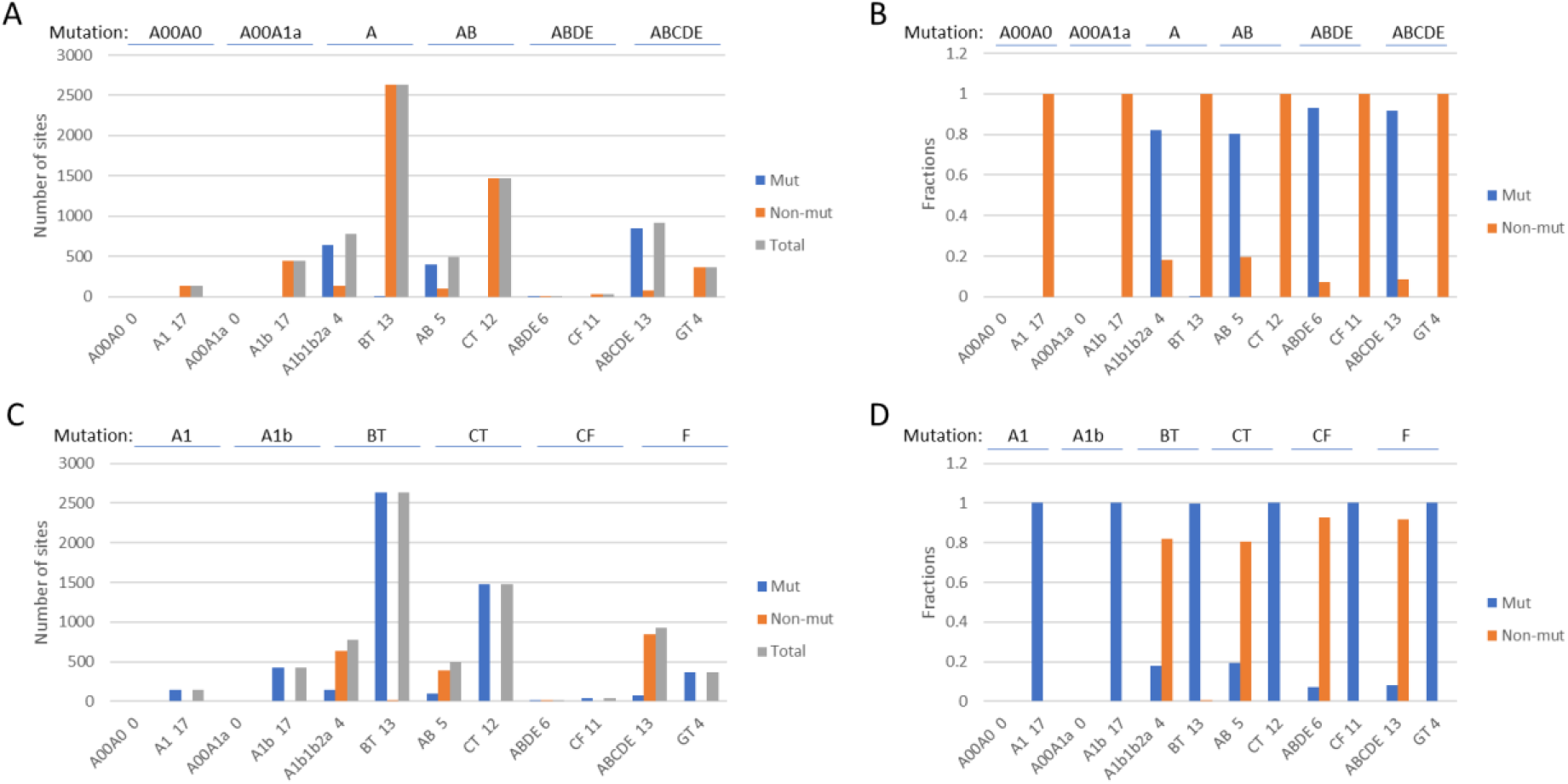
Expected mutation patterns only if the basal haplogroups in the Asia model are true. Shown are the number of mutated, non-mutated, and total informative sites in ancient DNAs under the Out of East Asia model (A) or the Out of Africa model (C). Also shown are the fractions of mutated and non-mutated sites in ancient DNAs under the Out of East Asia model (B) or the Out of Africa model (D). The number following the haplogroup name shows the number of ancient samples carrying the haplogroup.

Ancient haplogroups with some sites not mutated in the basal and terminal haplogroups to which they belong may not have persisted until today. Relative to the 36000 year old C1b K14 sample that had some sites not mutated in ABCDE-M89 and C-M130 basal haplogroups, the similarly aged C1a2 Sunghir SI, SIII, and SIV samples all had mutations in all sites informative to ABCDE and C-M130, and may better qualify as ancestors of some present day lineages. Regardless, ancient samples with only some, but not all, sites mutated for a basal haplogroup could serve to show that the basal haplogroups as inferred by studying present day samples did indeed exist.

The mutation pattern in the ancient Y chromosomes as revealed here confirms the expectation that ancient haplogroups should mutate in only a fraction of the sites that define a haplogroup they belonged to. Two observations here confirm the Asia model and invalidate the Africa model. First, only haplogroups specific to the Asia model showed the expected mutation pattern in ancient samples. Second, the genetic reality that a haplogroup, be it ancient or present, should not carry mutations found in basal haplogroups to which they do not belong is only met by the Asia model but not the Africa model. For example, I9133 A1b1b12a sample showed numerous shared alleles with BT, CT, CF, and F of the Africa model, to which it does not belong. However, it didn’t share any alleles with any haplogroups specific to the Asia model such as A00A0, to which it does not belong. We conclude that the Asia rooting of the Y chromosome phylogenetic tree has been independently confirmed by ancient DNAs.

## Materials and Methods

Whole genome sequencing data of ancient DNAs were downloaded from links provided by the previous publications. These ancient DNA samples were shown in Table 1. BWA 0.7.10 was used to aligned the fastq format to the human genome GRCh37 ^25^. The mapped reads were then filtered and sorted using SAMtools 0.1.19 and Picardtools 1.107 (http://picard.sourceforge.net) ^26^. The Picardtools 1.107 was also used to remove duplicates. The SNPs of genotypes were called using GATK-3.2-2 software with mapping quality setting at Q=30 ^27^. Haplogroups defining SNPs were identified using the ISOGG2018 Y chromosome tree (https://isogg.org) and the 1kGP dataset (http://www.intemationalgenome.org).

## Acknowledgments

Supported by the National Natural Science Foundation of China grant 81171880 (S.H.).

## Additional Information

### Author contributions

H.C. and Y.Z. performed data analysis. S.H. devised the project, analyzed the data, and wrote the manuscript. All authors edited the manuscript.

